# Encoding of vibrotactile stimuli by mechanoreceptors in rodent glabrous skin

**DOI:** 10.1101/2024.01.29.577766

**Authors:** Laura Medlock, Dhekra Al-Basha, Adel Halawa, Christopher Dedek, Stéphanie Ratté, Steven A. Prescott

## Abstract

Somatosensory coding in rodents has been mostly studied in the whisker system and hairy skin, whereas the function of low-threshold mechanoreceptors (LTMRs) in rodent glabrous skin has received scant attention, unlike in primates where glabrous skin has been the focus. The relative activation of different LTMR subtypes carries information about vibrotactile stimuli, as does the rate and temporal patterning of LTMR spikes. Rate coding depends on the probability of a spike occurring on each stimulus cycle (reliability) whereas temporal coding depends on the timing of spikes relative to the stimulus cycle (precision). Using *in vivo* extracellular recordings in rats and mice, we measured the reliability and precision of LTMR responses to different tactile stimuli including sustained pressure and vibration. Similar to other species, rodent LTMRs were separated into rapid-adapting (RA) or slow-adapting (SA) based on their response to sustained pressure. However, unlike the dichotomous frequency preference characteristic of RAI and RAII afferents in other species, rodent RAs fell along a continuum. Fitting generalized linear models (GLMs) to experimental data reproduced the reliability and precision of rodent RAs. The resulting model parameters highlight key mechanistic differences across the RA spectrum; specifically, the integration window of different RAs transitions from wide to narrow as tuning preferences across the population move from low to high frequencies. Our results show that rodent RAs can support both rate and temporal coding, but their heterogeneity suggests that co-activation patterns play a greater role in population coding than for dichotomously tuned primates RAs.

**Significance Statement:** Our sense of touch starts with activation of nerve fibres in the skin. Although response properties of various fibre types are well-established in other species (e.g. primates), quantitative characterization in rats and mice is limited. To fill this gap, we performed a comprehensive electrophysiological investigation into the coding properties of tactile fibres in rodent non-hairy skin and then simulated these fibres to explain differences in their responses. We show that rodent tactile fibres resemble those from other species, but that their heterogeneity at the population level may differ, with potentially important implications for encoding of touch. Simulations reveal intrinsic mechanisms that support this heterogeneity and provide a useful tool to explore somatosensation in rodents.

## Introduction

Processing of touch begins when low-threshold mechanoreceptors (LTMRs) in the skin convert mechanical stimuli into action potentials, or spikes, that are relayed to the CNS. Classic work classified LTMRs as slow-adapting (SA) or rapid-adapting (RA) based on responses to sustained pressure (1). Subsequent work, mostly in primate glabrous skin, subclassified SAs and RAs based on preferred frequencies. SAs spike repetitively during sustained pressure and are most sensitive to low-frequency vibrations (2–4) whereas RAs spike transiently during sustained pressure and are subdivided into RA1s, which are most sensitive to <50 Hz vibrations perceived as flutter, and RA2/PCs (Pacinians), which are most sensitive to >100 Hz vibrations perceived as continuous vibration (5). That said, the stimulus waveform is important (6) and RA2/PCs can respond to low-frequency pulse trains if the component pulses have sharp onsets (7), thus revealing the importance of disambiguating the frequency of repeating pulses from pulse kinetics.

Work in rats (8–13) and mice (14–19) has confirmed that rodent LTMRs resemble LTMRs in other species and has advanced our molecular understanding of somatosensation [for reviews see (20, 21)]. But the coding properties of rodent LTMRs have not been as thoroughly quantified as in primates (see above), cats (22, 23), or raccoons (24). The limited quantitative characterization of vibrotactile responses in rodents shows marked variability, especially in RA tuning (9, 16). Furthermore, despite emerging evidence of the importance of spike timing for tactile sensation (25–29), this has not yet been examined in rodent glabrous skin. Advances in genetic and optical techniques have furthered the use of rodent models for studying touch [for reviews see (21, 30)] in which glabrous skin, in particular, is stimulated to assess pain (e.g. von Frey and Hargreaves tests) (31–33). Therefore, a comprehensive investigation into the tactile coding properties of LTMRs in rodent glabrous skin is vital for our understanding of normal somatosensation and its pathological disruption.

An LTMR’s response to vibration is influenced by stimulus intensity (defined here as the peak-to-peak amplitude of force variations) and frequency. Previous studies suggest that stimulus intensity is rate-coded and frequency is temporally-coded (2, 4, 25, 34–37). Temporal coding depends on spike timing relative to stimulus phase (i.e. spike precision, phase-locking) whereas rate coding reflects the probability of spiking on each stimulus cycle (i.e. spike reliability, entrainment). Reliability and precision are both important for synchronization of spikes across neurons (38), and synchrony is important for sensation (39, 40). Firing rate will increase with stimulus frequency if LTMR spikes entrain 1:1 with stimulus cycles but firing intermittently (skipping irregularly) prevents this (41, 42). Examining the reliability and precision of LTMR responses is crucial for understanding somatosensory coding at a single-neuron level and, in turn, how neurons work together at the population level.

Using rats and mice, we quantified the reliability and precision of LTMR responses to hind paw stimulation *in vivo*. Clustering based on reliability, which is reflected in the frequency tuning curve, revealed that rodent RAs exhibit a continuum of frequency preferences. On the other hand, increasing stimulus frequency universally increased spike precision. We further confirmed that poorer precision at low frequencies was due to co-variation of the stimulus waveform with frequency during sinusoidal stimulation; applying trains of abrupt-onset pulses at low frequencies improved spike precision. GLMs fit to experimental data reproduced the reliability and precision of RA responses to vibration and sustained pressure. The resulting model parameters highlight key mechanistic differences supporting the spectrum of frequency preferences. Together, our results show that LTMRs in rodent glabrous skin resemble LTMRs from other species but their heterogeneity at the population level may differ, with potentially important implications for somatosensory coding.

## Materials & Methods

### *IN VIVO* EXPERIMENTS

All procedures were approved by the Animal Care Committee of The Hospital for Sick Children (protocols #53451 and #47072) and were conducted in accordance with guidelines from the Canadian Council on Animal Care. Adult male Sprague Dawley rats (200-350 g) were obtained from Charles River, Montreal. Male and female mice (10-12 weeks, 20-30 g) with a C57BL/6 background expressing channelrhodopsin-2 in sensory afferents were produced by crossing *Advillin^Cre/+^* mice (kindly provided by Fan Wang, Duke University) or *PV^Cre/+^* mice (JAX:017320) with Ai32 mice (JAX:024109), but optogenetic stimuli were not tested in this study. Rats and mice were anesthetized with 1.2 g/kg urethane i.p and received 10% top-ups as needed to block the pedal reflex. Laminectomies were performed to expose the L4 and L5 dorsal root ganglion (DRG) for single-unit extracellular recordings as previously described in rats (39) and mice (43). For rats, we used a multielectrode array with 16 contacts on 4 shanks (NeuroNexus, A4 type). For mice, we used a parylene-insulated tungsten microelectrode (A-M Systems, #573500). Signals were amplified, digitized at 40 kHz and high-pass filtered at 300 Hz using an OmniPlex Data Acquisition System (Plexon). Gentle brushing was used to search for LTMRs innervating the glabrous skin of the hind paw. Receptive fields were then mapped using von Frey filaments. Once a unit was identified, precise and reproducible stimuli were applied with a flat-tipped indenter (1 mm diameter) using a computer-controlled mechanical stimulator (Aurora Scientific, model 300C-I) positioned with a micromanipulator. Stimuli included sustained pressure steps (25-250 mN), sinusoidal vibration (2–400 Hz, 150–225 mN), and trains of brief pulses (10-50 pulses/second, 40% duty cycle, 150-225 mN) each applied for 1 second for each frequency-amplitude combination. All stimuli were controlled by a Power1401 A-D board and Signal v5 software (Cambridge Electronic Design). Stimulus timing was sent as a trigger to Plexon to align neural responses to stimuli. Single units were isolated using Offline Sorter v4 software (Plexon) and analyzed with MATLAB (MathWorks, R2021).

### SPIKE TRAIN ANALYSIS

Reliability and precision of LTMR responses to periodic tactile stimuli were measured using entrainment and temporal dispersion, respectively. Entrainment was calculated as

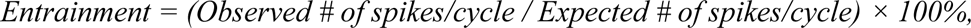

where the expected number of spikes per cycle is 1. We plotted entrainment against stimulus frequency to identify the preferred frequencies (tuning) of each neuron. Temporal dispersion is inversely related to spike timing precision and, for periodic stimuli, is also related to phase-locking. Phase-locking is typically expressed in radians or degrees, but here we convert it to absolute time by normalizing by the stimulus period using the formula,

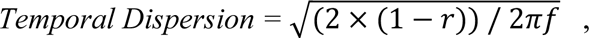

where *r* is vector strength, and *f* is stimulus frequency. Temporal dispersion is calculated based on the first spike in each cycle and is measured in milliseconds (ms). Conduction velocity for rat LTMRs was calculated by dividing the distance between the stimulation and recording sites (to heel: 0.12 m, mid-paw: 0.135 m, toe: 0.15 m) by the latency to the first spike from the onset of a strong, sustained pressure step.

### CLUSTER ANALYSIS

For an unbiased characterization of RA frequency preferences, K-means clustering was performed on RA tuning curves. For the clustering and principal component analysis (PCA), the features were individual stimulus frequency-intensity combinations (e.g. 10 Hz-150 mN) and the samples were the corresponding entrainment (e.g. 50%) for each feature. The optimal number of clusters for each K-value (between 2-5) was determined using the mean silhouette value. Z-scores were calculated and PCA was performed to visualize the high-dimensional clustering results. The first two PCs accounted for a total of 55.3% of the total variance in the data (PC 1: 34.7%, PC 2: 20.6%). Clustering was first performed on rat data and mouse data were subsequently assigned to existing clusters.

### GENERALIZED LINEAR MODELS (GLMs)

The GLM structure uses code modified from (44). Briefly, model parameters include a stimulus filter (*k*), a post-spike filter (*h*), and mean input (μ) that determines baseline firing rate. GLMs were fit to the response of individual RAs (n=20) to sinusoidal vibration across different combinations of frequency (5-400 Hz) and intensity (150, 175, 225 mN). To fit the GLM to experimental data, the stimulus and spike history (i.e. time-shifted spike trains) were each convolved with unique sets of basis functions and summed. Here, we used raised cosine basis functions to allow for fine temporal structure near the spike. The output of this convolution was then put through a nonlinear exponential function (*f*) to produce an instantaneous firing rate (λ) from which spike probabilities are calculated from a Poisson distribution. The negative log-likelihood function was minimized to find the optimal parameters (*k*, *h*, and μ). Once GLMs were fit, optimized filters were used to predict λ and spike probabilities in response to new stimuli. If the GLM generated a probability greater than a uniformly distributed random number between 0.001 and 1 (*rand*, MATLAB), a spike occurred, and the post-spike filter was applied. To validate each model, we tested its response to additional experimental data that were not used for training (5-400 Hz at 200 mN). GLMs were also validated against responses to sustained pressure steps.

The stimulus filter for each model LTMR was 50 ms and the number of basis functions ranged from 10 to 20. Using a grid search method, we optimized the number of basis functions for the GLM for each LTMR by minimizing the sum of the error between model and experimental reliability and precision across all trials (i.e. all frequency-intensity combinations). The length and number of basis functions for the post-spike filter were held constant at 85 ms and 10, respectively, as variations in these hyperparameters had little to no effect on model error.

### FILTER ANALYSIS

To analyze the fitted stimulus and post-spike filters for each GLM, we measured area under the curve (AUC). AUC was calculated using trapezoidal numerical integration (*trapz* function, MATLAB) for the entire filter, as well as its positive (+) and negative (−) components. A power spectrum analysis was performed for each k-filter using a discrete Fourier transform (*fft* function, MATLAB) which was converted to a double-sided power spectrum (P2) using, P2 = | fft(k) / L |, where L is filter length of 50 ms. The average (± SD) single-sided power spectrum for k-filters was plotted across different clusters. The width of the post-spike filter was also measured from time 0 to the time the filter first passed zero on the y-axis. This post-spike filter width is analogous to the neuron’s refractory period following a spike.

### MUTUAL INFORMATION ANALYSIS

A lower bound on mutual information was calculated by subtracting the entropy of the conditional distribution, P(**S**|**R**), from the entropy of the uniform prior distribution, P(**S**), where **R** is the set of response vectors and **S** is the set of stimulus labels (8 amplitudes x 16 frequencies; 7 bits of entropy). P(**S**|**R**) was approximated by a decoder (random forest) which was trained to classify both stimulus frequency and amplitude. The input to the decoder, r_i_ *ϵ* **R**, was a feature mapped representation of LTMR activity over 100 milliseconds. For single neurons, the feature map involved discretizing a sequence of inter-spike intervals into 10 bins, with edges logarithmically spaced between 1 and 100 ms, and using the counts in each to form a 10-dimensional input vector. Feature mapping for population activity was accomplished by summing the feature vectors of each neuron in that population. This mapping both improved classification performance and standardized the dimensionality of the input. Following training, the change in entropy between P(**S**) and P(**S|R**= r_i_) was computed for each test example (*i*), and these values were averaged together to produce the final measure of information between **S** and **R**. A new decoder was initialized and trained for each of the 3800 {**S, R**} pairs: 2 groups (heterogeneous vs. homogeneous; see below) x 100 permutations x 19 populations.

More homogenous or more heterogeneous populations of LTMRs were created to analyze population-level coding. To control for the sum of individual neuron informativeness within each type of population (homogeneous vs heterogeneous), LTMRs were split into eight groups of two or three neurons with similar levels of mutual information per neuron. A neuron is selected from each group to create a population of N neurons. Each population type starts with the same first neuron but all subsequently added neurons are chosen from randomly ordered groups. Neurons were chosen from each group based on their similarity in frequency tuning to the first neuron, as defined by their PC2 value. Specifically, homogenous populations were created by choosing neurons with similar PC2 values and heterogeneous populations were created by choosing neurons with dissimilar PC2 values. This process was repeated 100 times per population size (i.e. 100 permutations; see **Figure 8C**).

### STATISTICAL ANALYSIS

All statistical analysis was performed using GraphPad Prism 9.5.1 (GraphPad Software, LLC). Parametric tests were used when data were normally distributed (according to the Shapiro-Wilk test). The comparison of two groups was done using an unpaired t-test with Welch’s correction. When comparing more than two groups, a one-way ANOVA was performed, followed by post hoc Tukey’s tests when appropriate. Pearson’s correlation coefficient (*r*) and p-values are stated for all correlations. R-squared (R^2^) and p-values are reported for all linear regressions. Chi-square (χ^2^) and p-values are reported when comparing proportions.

### CODE AVAILABILITY

Code for models will be available on ModelDB upon publication.

## Results

### Rodent LTMRs respond heterogeneously to tactile stimuli

To characterize the responses of rodent LTMRs to tactile stimulation, we stimulated the hind paw of rats and mice with various tactile stimuli while recording from somata in the DRG. Similar to other species, rat LTMRs were classified as slow-adapting (SA, n=4) or rapid-adapting (RA, n=31) if they spiked repetitively or transiently, respectively, to steps of sustained pressure (**Fig. 1A**). In response to sinusoidal vibration, SAs consistently entrained best at low frequencies (<50 Hz), often firing >1 spike per stimulus cycle (**Fig. 1B**, green), whereas RAs were more heterogeneous (**Fig. 1B**, i-iv). Some RAs were tuned to low frequencies (<50 Hz; example ***i***), others were broadly tuned (examples ***ii*** and ***iii***), and some were tuned to high frequencies (≥50 Hz; example ***iv***). Whereas the frequency-sensitivity of entrainment varied across RAs, temporal dispersion consistently decreased with increasing frequency (**Fig. 1B**, bottom row). Similar response properties were observed in mouse RAs (**Fig. 2**). We proceeded to characterize RA responses in more detail.

**Figure 1–.**
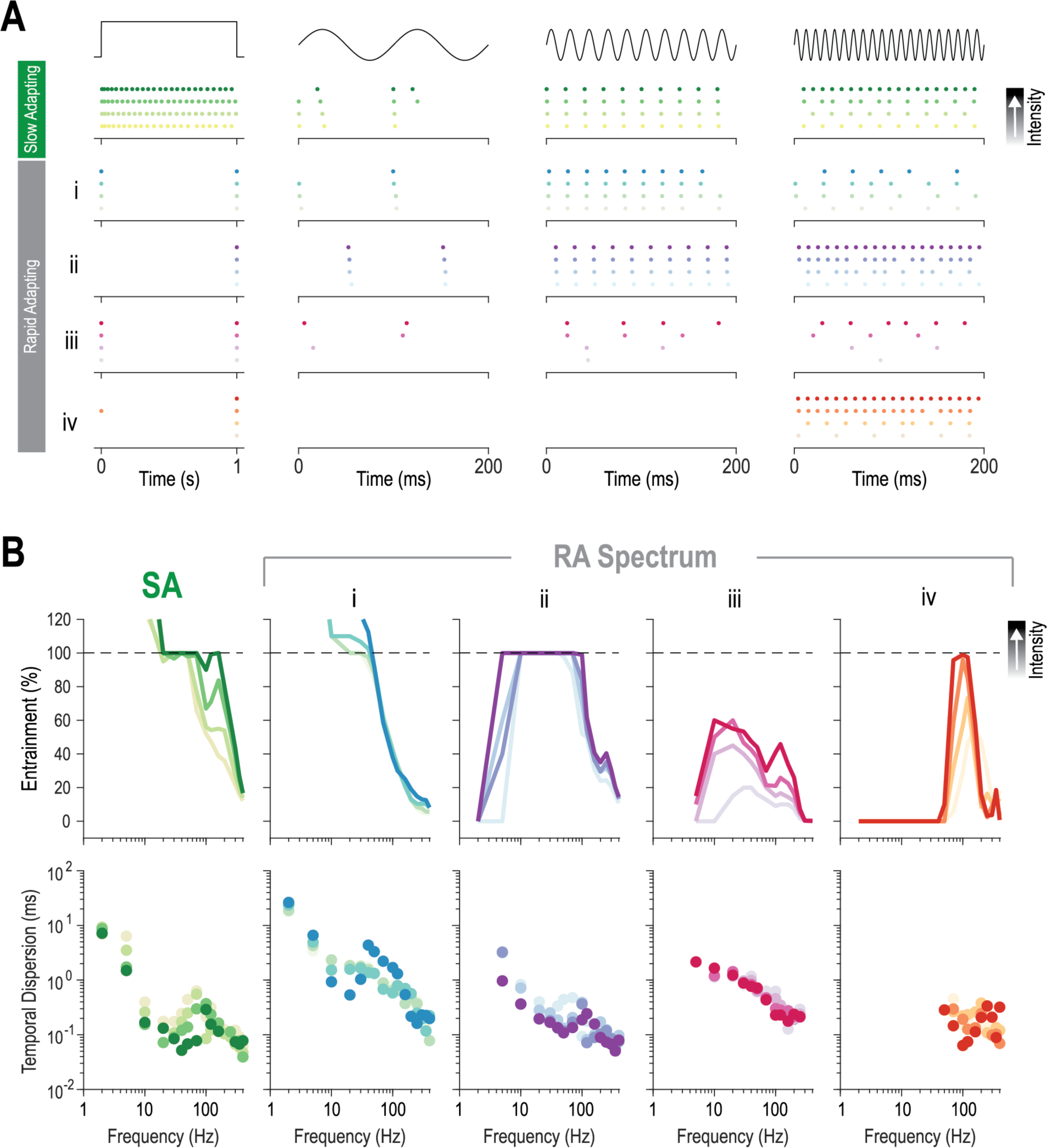
Heterogeneity in rat LTMR responses. **(A)** Sample responses to sustained pressure (left) and sinusoidal vibration at 10, 50, and 100 Hz from an SA and four RAs (*i-iv*) for four stimulus intensities (color density). Rasters depict each spike as a dot. **(B)** Entrainment (top) and temporal dispersion (bottom) as a function of stimulus frequency for units depicted in **A**. Dashed line at 100% entrainment corresponds to one spike per stimulus cycle. The four RAs shown here are representative neurons from the four clusters in Figure 3.

**Figure 2–.**
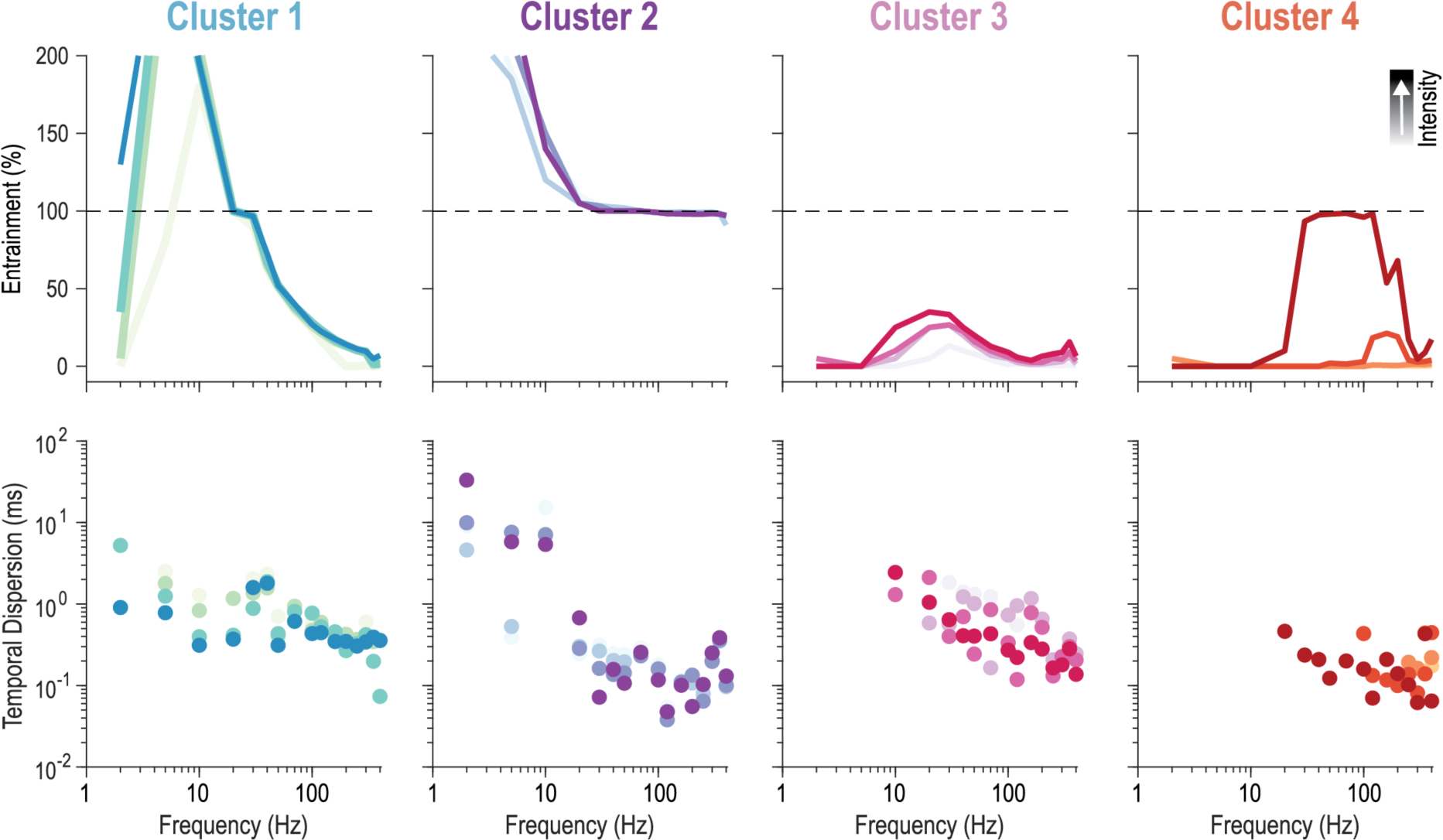
Heterogeneity in mouse RA responses. Entrainment (top) and temporal dispersion (bottom) for four stimulus intensities (color density) as a function of stimulus frequency for four RAs (one from each cluster from Figure 3).

### RAs exhibit a continuum of frequency preferences

To classify rat RAs based on their response to vibration, we applied K-means clustering to their tuning curves (**Fig. 3**). Clustering based on individual stimulus intensities (**Fig. 3A**) revealed three dominant clusters (filled circles) and a subset of neurons that changed clusters depending on stimulus intensity (open circles). Clustering based on all intensities (**Fig. 3B**) revealed four clusters, including three (Clusters 1, 2, 4) similar to those in panel A and a fourth (Cluster 3) representing neurons whose membership in the other three clusters varied with stimulus intensity. Mouse RAs (n=7; **Fig. 3B**, triangles) fell within clusters of rat RAs, and were assigned accordingly. Sample tuning curves for an RA from each cluster are shown in **Fig. 1B** (for rat) and **Fig. 2** (for mice). PC1 captures tuning curve width (more positive PC1 values indicate broader tuning) and PC2 captures frequency preference (more positive PC2 values indicate preference for lower frequencies) according to PC loading analysis (bottom right inset in **Fig. 3B**). Rather than revealing clearly separable clusters, this analysis suggests that neurons fall along a continuum, especially when considered in a single dimension (i.e. tuning breadth *or* preferred frequency).

**Figure 3–.**
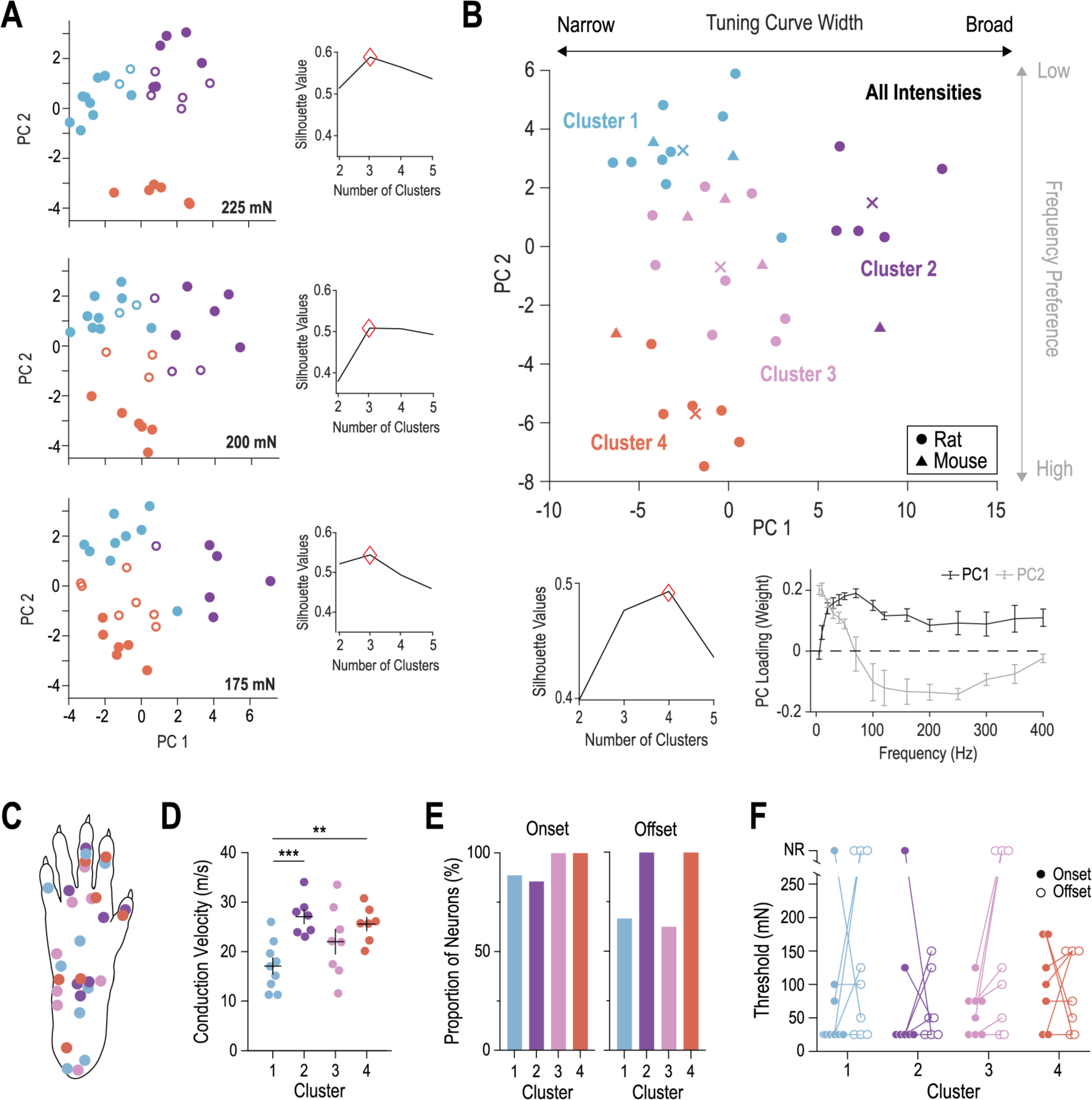
Rat RAs do not fall into discrete subtypes but, instead, exhibit a continuum of frequency preferences. **(A)** K-means clustering of RAs based on individual stimulus intensities (175, 200, 225 mN) suggested three clusters (blue, purple, orange) but certain neurons (open circles) did not fall consistently into the same cluster, and instead varied depending on stimulus intensity. Optimal number of clusters was determined from the peak in silhouette values (red diamonds, inset). **(B)** Clustering based on all stimulus intensities suggested four clusters (see silhouette values on bottom left inset). Centroids are marked with x. Cluster 3 comprises RAs depicted as open circles in panel **A**. Mouse RAs (triangles) were assigned to appropriate rat clusters after projecting data on PC1 and PC2, which reflect tuning curve width and preferred frequency, respectively, according to PC loading (bottom right inset). Mean (± SEM) PC load is calculated from all stimulus intensities at each stimulus frequency (see Methods). **(C)** Receptive field centers for individual RAs. **(D)** Conduction velocity (CV) differed significantly between some of the clusters (F_3,27_=6.062, p=0.003; one-way ANOVA, Tukey’s post-hoc). Specifically, CV of Cluster 1 (17.08 ± 1.65 m/s) was higher than Cluster 2 (27.06 ± 1.44 m/s, ***p<0.001) and Cluster 4 (25.56 ± 1.35 m/s, **p=0.003) but not Cluster 3 (22.01 ± 2.50 m/s). **(E)** Proportion of neurons spiking transiently at the onset or offset of sustained pressure steps did not differ significantly across clusters (onset: χ^2^=2.0701, p=0.5580; offset: χ^2^=6.1742, p=0.1034) **(F)** Threshold for onset and offset spikes often differed within a given RA, but not with any pattern that was consistent across clusters.

Further comparison of different RA properties revealed that clustering was independent of the receptive field location (**Fig. 3C**) and there was little difference in conduction velocity (**Fig. 3D**), presence of onset/offset spikes (**Fig. 3E**), or the threshold for onset/offset spikes (**Fig. 3F**) across the four clusters. These findings suggest that our clusters do not represent distinct subpopulations of RAs with unique anatomical RF locations or electrophysiological properties but, instead, represent a heterogeneous population with a continuum of frequency preferences.

### Models reproduce RA responses to vibration and pressure

To further study RA properties, we fit GLMs to responses to a subset of vibrotactile stimuli (**Fig. 4A**). All successfully fitted models (n=20) reproduced responses to vibrotactile stimuli not included in the training data set (**Fig. 4B**). The models for RAs illustrated in **Figure 1i-iv** are shown in **Figure 4Bi-iv**, respectively. The entrainment and temporal dispersion of simulated (predicted) responses did not deviate significantly from experimental data (see insets). Like experimental data, model RAs exhibited heterogeneous tuning curves and temporal dispersion decreased with increasing stimulus frequency (**Fig. 4B**).

**Figure 4–.**
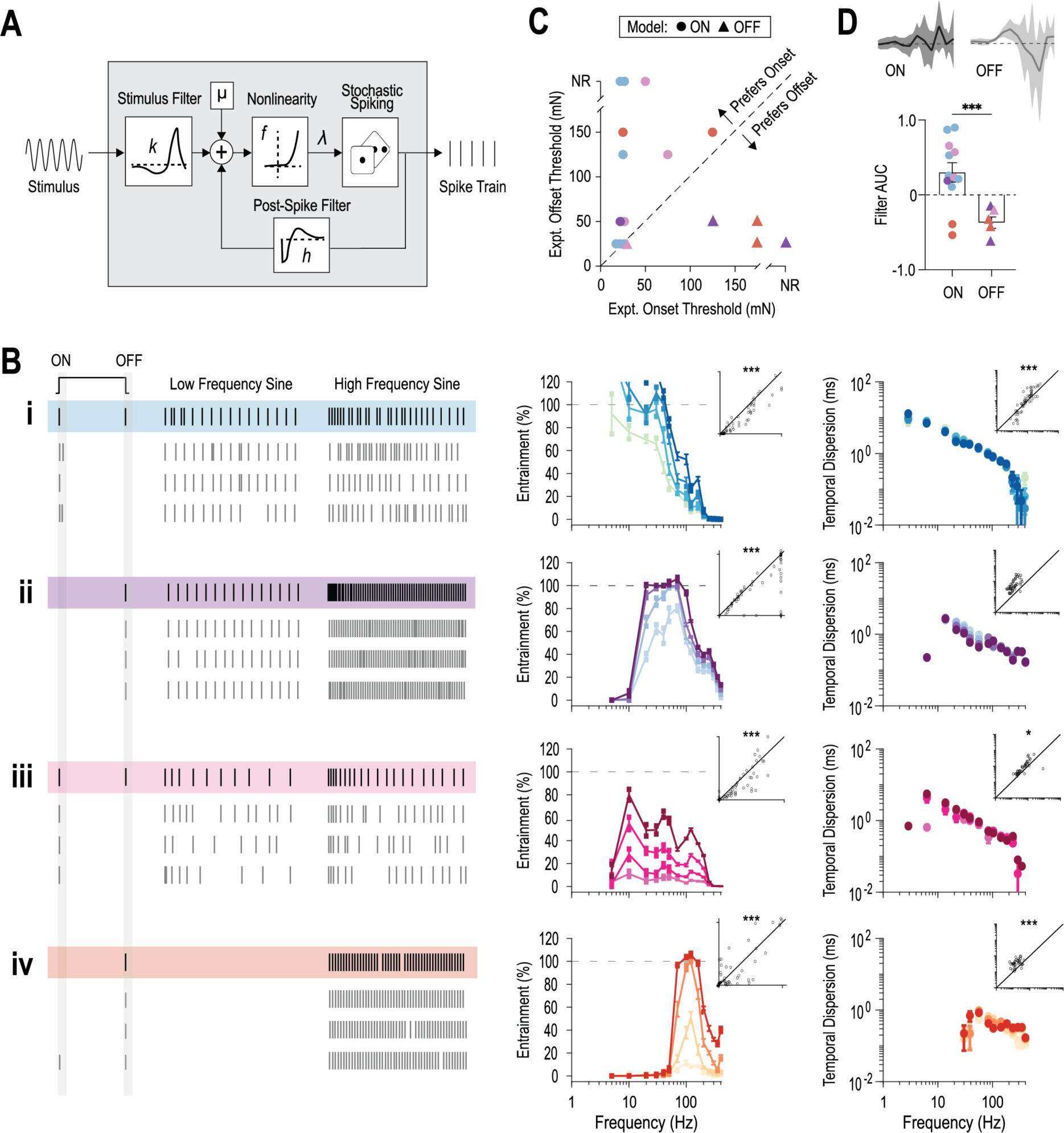
Fitted GLMs quantitatively reproduce RA response properties. **(A)** Schematic of GLM components modified from (44). **(B)** Sample spike trains (left) and corresponding entrainment (middle) and temporal dispersion (right) for GLMs fitted to units *i-iv* in Fig. 1B. Experimentally observed rasters (colored background) are shown for comparison against three spike trains simulated from the corresponding GLM. Insets show entrainment and temporal dispersion of simulated (predicted, y-axis) responses plotted against experimental (observed, x-axis) responses. Data points, each representing a different frequency-intensity combination, tend to fall along the diagonal indicating equivalence. Correlations for reliability [*i*: *r*=0.91, ***p=1.47e-23; *ii*: *r*=0.67, ***p=6.83e-09; *iii*: *r*=0.87, ***p=7.92e-20; *iv*: *r*=0.83, ***p=3.55e-16]. Correlations for temporal dispersion [*i*: *r*=0.80, ***p=3.12e-14; *ii*: *r*=-0.14, p=0.288; *iii*: r=0.26, *p=0.0440; *iv*: r=0.69, ***p=8.53e-10]. **(C)** Threshold for onset and offset spikes in model RAs. Models divided into two groups, ON (circles) or OFF (triangles), based on spiking at onset or offset to sustained pressure. There was no consistent alignment between ON/OFF model grouping and Clusters 1-4 (from Fig. 2) except for Cluster 1 RAs whose corresponding models were all ON. Deviation above or below diagonal (dashed line) indicates a preference (lower threshold) for onset or offset spiking, respectively, in experimental data from corresponding RAs. Models accurately predict this preference (i.e. ON models had lower experimental onset threshold and OFF models had lower offset threshold). **(D)** Average stimulus filter for ON (black) and OFF (grey) models (shading=SEM). OFF models had a more negative filter (quantified as net area under the curve, AUC) compared to ON models (***p=0.0004; Welch’s t-test), consistent with their preference for decreases in stimulus force (i.e. stimulus offset).

To further validate models, we also simulated responses to sustained pressure. Models responded at stimulus onset or offset (**Fig. 4C**), and are labelled ON or OFF, respectively; OFF models also responded at the onset of high-amplitude steps. Experimental data confirmed that neurons fitted with OFF models had a lower threshold for offset spikes than for onset spikes (i.e. prefer stimulus offset) whereas the opposite was true for neurons fitted with ON models. Since responses to sustained pressure were not included in the training data set, reliable reproduction of these responses demonstrate that our models can accurately predict a neuron’s preference for stimulus onset or offset. Indeed, analysis of model stimulus filters revealed that OFF neurons have a more negative filter (quantified as area under the curve) compared to ON neurons (**Fig. 4D**; p=0.0004) which supports their preference for downward deflections in stimulus position (i.e. stimulus offset). Together, these simulations confirm that our models accurately reproduce RA responses to diverse tactile stimuli.

### Stimulus filters capture the continuum of frequency preferences in RAs

Following model validation, we analyzed RA filters to explore the mechanisms underlying their heterogeneous frequency tuning. We hypothesized that the stimulus filter, which describes a neuron’s preferred stimulus feature, would parallel the preferred frequency as ascertained from the entrainment tuning curves (see **Fig. 1**). As predicted, ordering of RA stimulus filters according to PC2 value revealed a continuum of filter widths (**Fig. 5A**). Specifically, stimulus filters transitioned from wide to narrow as tuning preference across the RA population moved from low to high frequencies (i.e. from positive to negative PC2 values). To quantify this change, we calculated the power spectrum for individual RA filters and found the average within each cluster (**Fig. 5B**). Cluster 1 filters had peak power at low frequencies (<50 Hz), Cluster 2 and 3 filters had broader, bimodal power spectra, and Cluster 4 filters had peak power at high frequencies (>100 Hz). This gradual transition of peak filter power from low to high frequencies further supports that RAs form a frequency preference continuum.

**Figure 5–.**
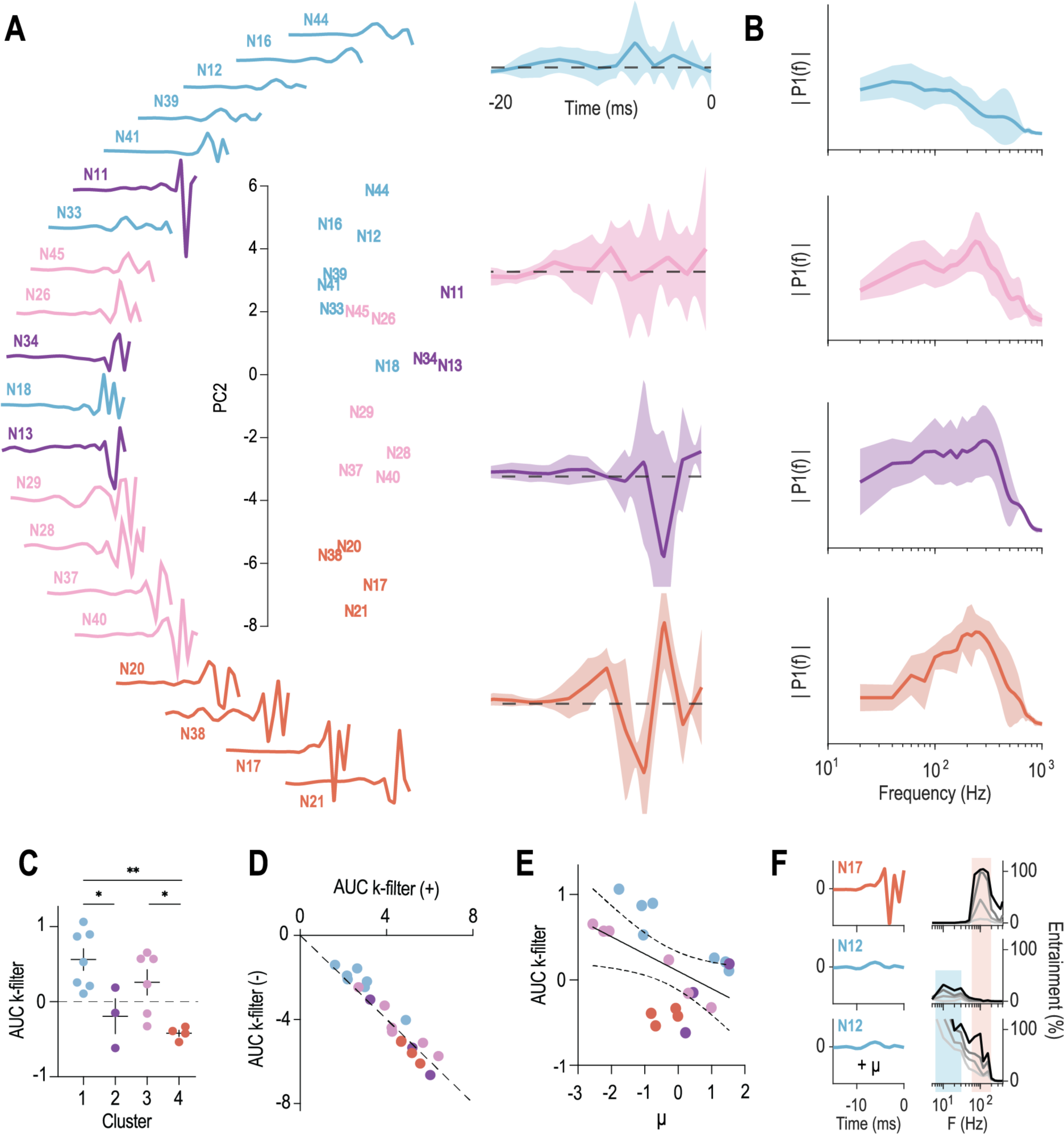
Stimulus filters of model RAs capture their spectrum of frequency preferences. **(A)** Stimulus filters of RA models are shown aligned from smallest to largest PC2 value. Filters transition from wide to narrow as PC2 values move from positive to negative (i.e. frequency preference shifts from low to high). Mean stimulus filters (right) for each cluster (shading=SEM). Positive and negative components of each filter are above and below the dashed line at zero. **(B)** Average power spectrum for each cluster. Peak power shifts from low frequency for Cluster 1 filters to high frequency for Cluster 4 filters. Shading=SEM. **(C)** Net AUC of the stimulus filter shifted from positive to negative across Clusters 1 to 4 (i.e. as frequency preference moved from low to high). Stimulus filter AUC differed significantly between clusters (F_3,16_=7.402, p=0.0025; one-way ANOVA). Specifically, Cluster 1 units had more positive AUC than Cluster 2 (*p=0.0368) or Cluster 4 (**p=0.0026) and Cluster 3 units had more positive AUC than Cluster 4 (*p=0.0460; Tukey’s post hoc tests). **(D)** Positive (+) versus negative (−) component of stimulus filter AUC. Data points represent stimulus filters for individual units and fall along the diagonal (dashed line) indicating a balance between + and – AUC components. Generally, neurons preferring low frequency (e.g. Cluster 1) had smaller stimulus filters (small + component balanced by small – component) than neurons preferring high frequency (e.g. Cluster 4; large + component balanced by large – component). **(E)** Negative correlation between stimulus filter AUC and μ (R^2^=0.2483, p=0.025). **(F)** Swapping a narrow stimulus filter (orange; N17) and wide stimulus filter (blue; N12) transitions the frequency preference of the neuron from high (orange band) to low (blue band). Swapping the μ for N17 and N12 increased entrainment to ≥100% (see relationship in **E**).

To quantify additional differences, we analyzed the area under the curve (AUC). Stimulus filter AUC shifted from net positive to net negative as frequency preference shifted from low to high (i.e. from Clusters 1 to 4) (**Fig. 5C**). Decomposing stimulus filters into their positive (+) and negative (−) components showed that neurons with low-frequency preferences have small + components balanced by small – components, whereas neurons with high-frequency preferences have large + components balanced by large – components (**Fig. 5D**). Small imbalances account for the net AUC differences observed in Figure 5C. Together these results quantify the shift in RA stimulus filters from being smaller, more positive filters for neurons with low-frequency preferences to bigger, more negative filters for neurons with higher-frequency preferences. There was also a strong negative correlation between stimulus filter net AUC and μ (**Fig. 5E**; R^2^=0.2483, p=0.025), which suggests that neurons with more strongly negative stimulus filters require a larger (more positive) baseline current to help promote spiking. Finally, to demonstrate the role of the stimulus filter in determining a neuron’s frequency preference, we swapped a narrow (N17, orange) and wide (N12, blue) stimulus filter which caused the model to transition from high to low frequency preference (**Fig. 5F**; orange to blue band in entrainment plots). Since we found a relationship between stimulus filter and μ (see **Fig. 5E**), we also swapped the μ for N17 and N12 to help promote spiking and increase entrainment to ≥100% (bottom panel of **Fig. 5F**). In summary, analysis of model parameters revealed that cell-to-cell variability in the stimulus filter supports heterogeneity in RA tuning and further suggests that heterogeneity in stimulus filter width (i.e. length of the integration time window) underlies the continuum of RA frequency preferences.

### Post-spike filters capture the ability of RAs to spike repetitively

To separate the effects of waveform kinetics from repetition rate on LTMR responses (since the maximum rate of change of force increases with sinusoid frequency), we also tested square pulses repeated at 10-50 Hz. Unlike their response to low frequency sinusoidal vibration, most RAs entrained at ≥100% to low-frequency pulse trains and did so with low temporal dispersion. Specifically, plotting entrainment and temporal dispersion for sinusoids vs pulse trains for the equivalent frequencies and intensities shows that entrainment was significantly higher (**Fig. 6A**) and temporal dispersion was significantly lower (**Fig. 6B**) for pulse trains. Entrainment to pulses dropped in nearly all neurons as pulse rate was increased, though we were limited to testing ≤50 Hz for technical reasons. The same trend of decreasing entrainment was evident with increasing sinusoid frequency in cluster 1 neurons, whereas cluster 2 and 3 neurons exhibited increasing entrainment while cluster 4 neurons did not spike. Whereas the latter trends (to sinusoids) mostly reflect differences in frequency tuning across clusters (see Fig. 5B), the former trend (to pulse trains) uniquely reflects the refractoriness of a neuron (i.e. how quickly it recovers from the last pulse in order to respond to the next pulse). Refractoriness is reflected in the width of the negative component of the post-spike filter.

**Figure 6–.**
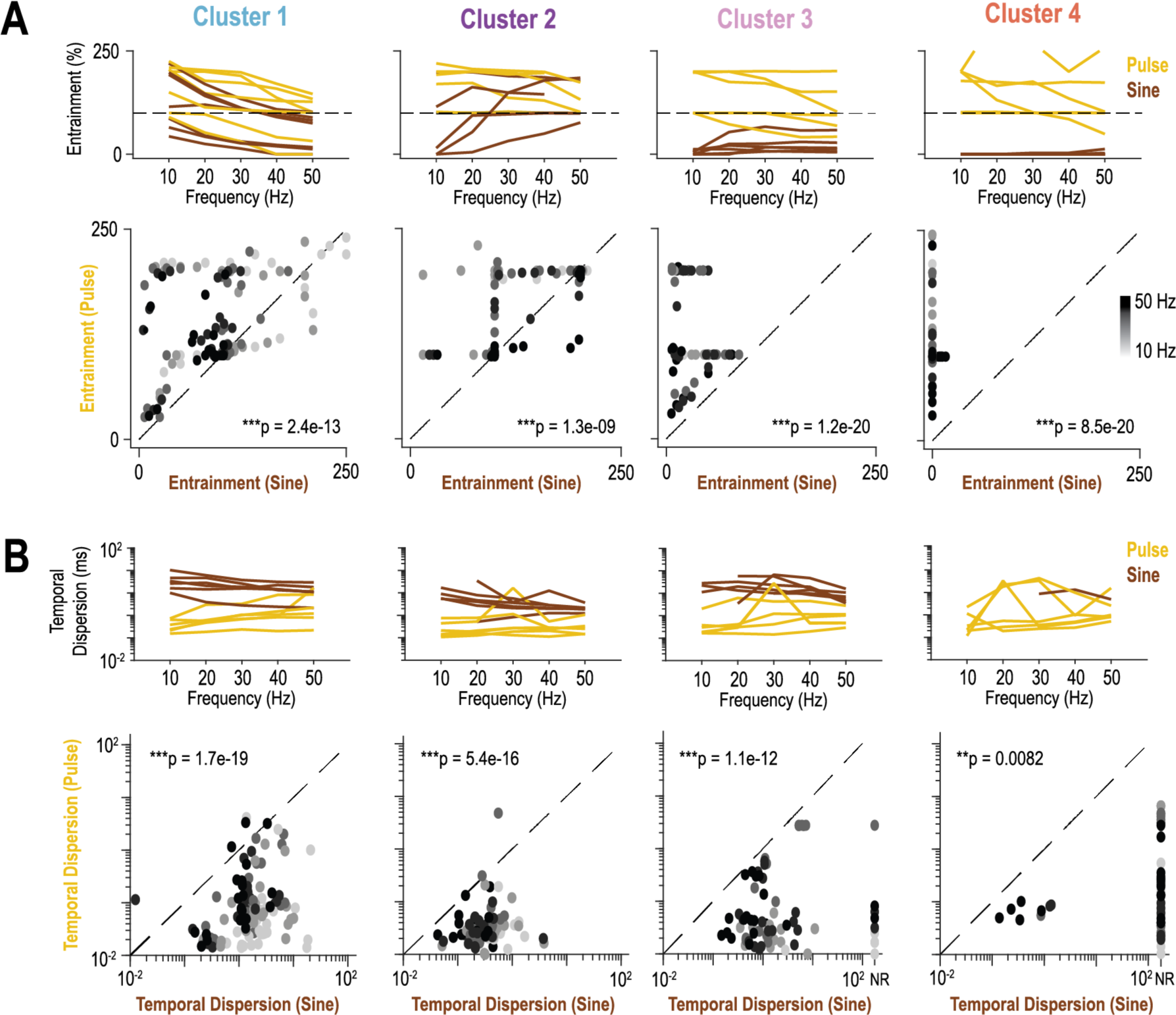
Response of RAs to pulse trains. (**A)** Entrainment is higher during pulse trains (yellow) than sinusoidal stimulation (brown) at equivalent frequencies (C1: ***p=2.43e-13; C2: ***p=1.28e-09; C3: ***p=1.20e-20; C4: ***p=8.46e-20; Wilcoxon signed rank tests). **(B)** Temporal dispersion is lower during pulse trains than sinusoidal stimulation (C1: ***p=1.69e-19; C2: ***p=5.36e-16; C3: ***p=1.11e-12; C4: **p=0.00815; Wilcoxon signed rank tests). **(A&B, top)** Each line depicts the mean entrainment (A) or temporal dispersion (B) for a single unit averaged across four stimulus intensities. Dashed line represents 100% entrainment. **(A&B, bottom)** Each dot represents reliability (**A**) or precision (**B**) of an individual neuron’s response to pulse and sine stimuli at the same frequency (10-50Hz) and amplitude. Data points sit above (**A**) or below (**B**) the diagonal (dashed line) indicating significantly higher reliability and significantly lower temporal dispersion, respectively (see p-values on plots; Wilcoxon signed rank test), to square pulses than to smooth sinusoids.

Unlike the stimulus filter, the post-spike filter of fitted GLMs did not change in a consistent manner across the frequency preference spectrum (**Fig. 7**). Ordering of post-spike filters according to PC2 value revealed heterogeneity in this model parameter across the RA population (**Fig. 7A**), but no significant difference in post-spike filter width (**Fig. 7B**; p=0.463) or AUC (**Fig. 7C**; – component: p=0.257, + component: p=0.404) between clusters. To explore the role of the post-spike filter on RA response properties, we simulated responses to square pulses repeated at different frequencies (**Fig. 7D**). Models entrained 1:1 with low frequency (≤50 Hz) pulse trains. As frequency was increased, models could not maintain 1:1 entrainment as the interpulse interval dropped below their post-spike filter width. For example, N12 (lighter blue) had a filter width of ∼20 ms and therefore could not entrain 1:1 with pulse trains >50 Hz (i.e. cycle time <20 ms) whereas N39 (darker blue) had a shorter filter width of ∼10 ms and could entrain 1:1 at ≥ 100 Hz. Both neurons had low entrainment at 200 Hz because the cycle length (5 ms) was shorter than their post-spike filters.

**Figure 7–.**
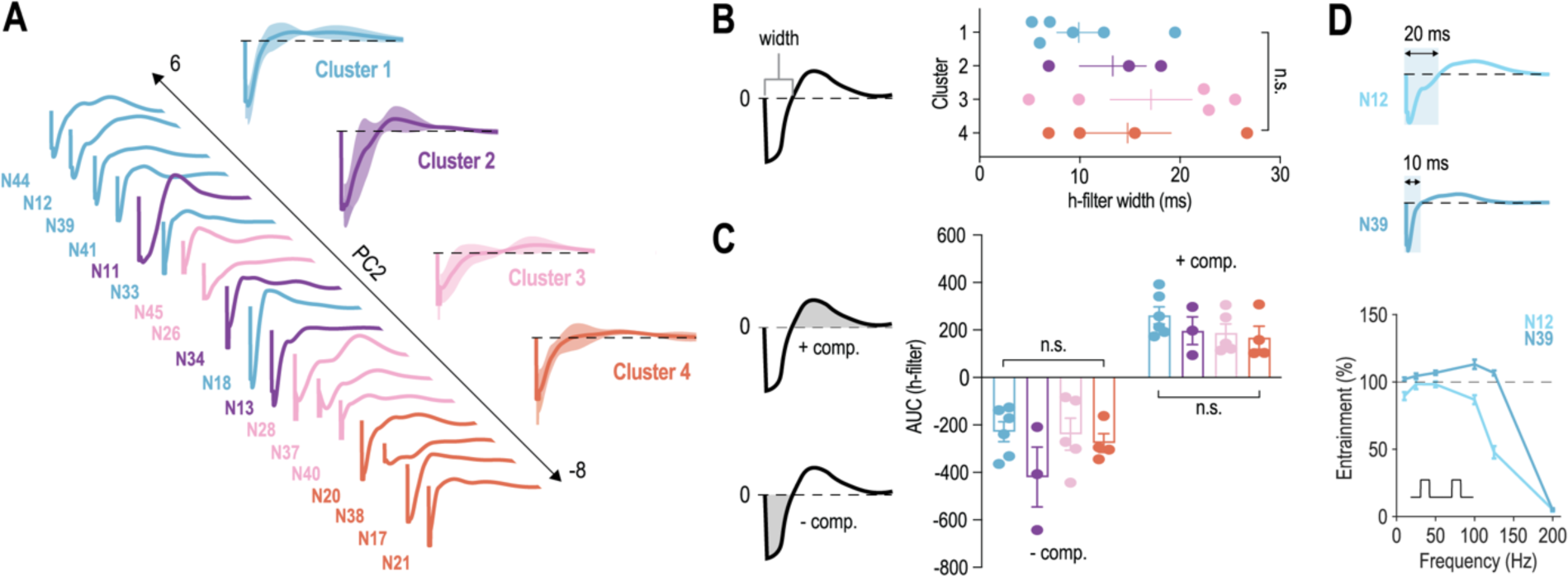
Post-spike filters of model RAs capture the ability to spike repetitively rather than frequency preference. **(A)** Post-spike filters aligned by the neuron’s PC2 value, from low to high frequency preference, show no consistent difference across clusters (heterogeneity is similar within and between clusters). Average post-spike filters (±SEM, shading) are similar across Clusters 1 to 4. **(B)** Width of post-spike filters did not differ significantly across clusters (F_3,14_=0.9603, p=0.4628; one-way ANOVA). **(C)** Positive (+) and negative (−) AUC components of post-spike filter did not differ significantly across clusters (+ AUC: F_3,14_=1.043, p=0.4042; – AUC: F_3,14_=0.502, p=0.2572; one-way ANOVAs). **(D)** Width of post-spike filter impacts model neuron’s ability to spike repetitively to pulse trains. N12 has a longer post-spike filter width (∼20 ms, light blue) than N39 (∼10 ms, darker blue) and therefore cannot 100% entrain to pulse trains above 50 Hz (cycle length = 20 ms). N39 continues to 100% entrain at ≥ 100 Hz (cycle length = 10 ms). Neither neuron spikes reliability at 200 Hz because the cycle length (5 ms) is shorter than their post-spike filter widths.

### Heterogenous tuning improves the efficiency of population-level coding

Finally, to probe the functional consequences of heterogeneity in LTMR tuning, we compared the amount of mutual information on stimulus identity from homogeneous and heterogeneous populations of LTMRs. We hypothesized that, all else being equal, a heterogeneous population of RAs conveys more information about a stimulus than a homogeneous population because the individual neuron responses carrying that information are less redundant in the former case (**Fig. 8A**). In other words, if several neurons are co-activated by various stimuli, together those neurons will convey more information if their tuning is more different (less overlapping) than if their tuning is more similar (more overlapping). To test this, populations were created that either favoured neurons with similar frequency preferences (homogeneous populations) or dissimilar ones (heterogeneous populations) (**Fig. 8C**). As predicted, our analysis revealed that heterogenous RA populations convey significantly more information about stimulus identity than homogenous populations when populations contained between six and seventeen neurons (**Fig. 8B**; *p<0.001, #p<0.01). These results demonstrate that heterogeneity in LTMR tuning allows for efficient population-level coding.

**Figure 8–.**
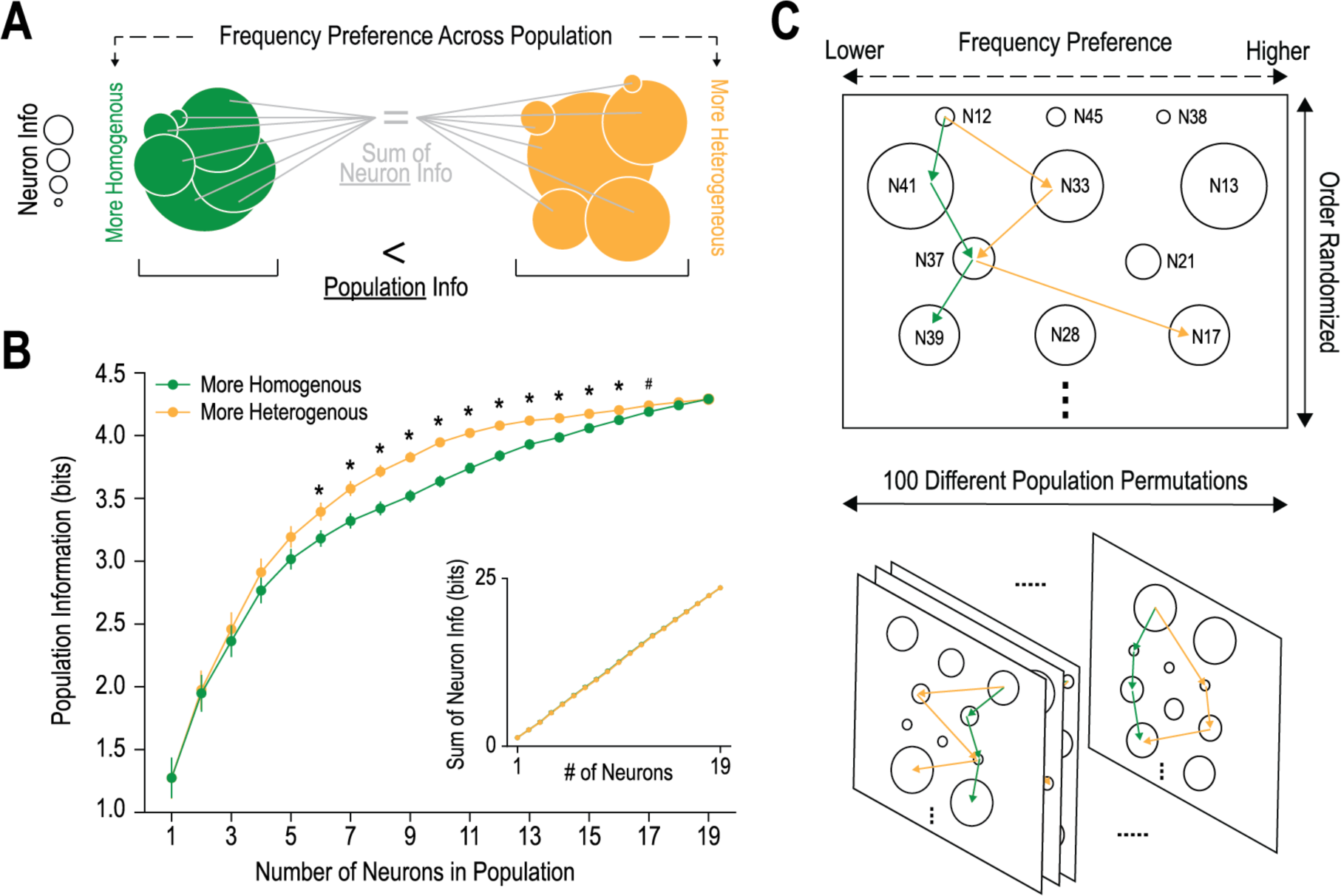
Heterogeneous tuning improves the efficiency of population-level coding. **(A)** Schematic outlining hypothesis that populations comprising an equal number of neurons with comparable informativeness per neuron (represented by circle area) will have greater mutual information on stimulus identity if constituent neurons have heterogeneous frequency preferences (yellow) than if they are more homogeneously tuned (green). Venn diagrams depict amount of overlapping information for each population type. The sum of neuron informativeness (circle area) must be controlled for because it is the upper bound on mutual information in a population code. **(B)** Heterogeneous populations (yellow) contain significantly more mutual information than homogeneous populations (green) (*p<0.001, ^#^p<0.01; mean±2 SEM, n=100 permutations; t-tests with Bonferroni correction). Inset: The sum of neuron informativeness is equivalent for each population type. **(C)** Overview of procedure for sampling neurons that are more similar (homogenous) or different (heterogeneous) based on frequency preference (PC2 value), when controlling for individual neurons’ informativeness (see **B**, inset).

## Discussion

Using in vivo single-unit recordings of LTMRs innervating glabrous skin, we have thoroughly characterized rodent LTMR coding properties. We found that rodent RAs respond heterogeneously to sustained and vibratory force (**Figs. 1 & 2**), forming a continuum of frequency preferences across the population (**Fig. 3**). Fitted GLMs reproduced experimentally observed response characteristics (**Fig. 4**) and linked the continuum in frequency preference to a continuum in stimulus filter shape (**Fig. 5**). The post-spike filter did not correlate with the stimulus filter but helps explain entrainment at high frequencies (**Fig. 7**). We also demonstrate that heterogeneity in tuning can improve efficiency of population-level coding (**Fig. 8**).

The sensing of skin vibrations by LTMRs is vital for the perception of object properties (e.g. texture) (45–47) and for sensorimotor control (e.g. slip detection) (2, 48, 49). In humans, monkeys, and cats, RA1s are most sensitive to ∼30 Hz whereas RA2/PCs are most sensitive to ∼250 Hz (2, 22, 49, 50). In contrast to this seemingly clear dichotomy, we found that rodent RAs are heterogenous in their frequency preferences and sometimes have very broad tuning curves (**Fig. 3**). Previous studies in raccoons and cats also showed variability in RA tuning (22, 51, 52) and this is also evident in recent mouse data (16). Conversely, RA tuning curves in monkey glabrous skin appear less heterogeneous (2, 53), but are nonetheless quite wide, such that RA1s and RA2/PCs are co-activated by natural stimuli (54). Primate RAs are essential for object manipulation and are localized to skin areas necessary for gripping (55) whereas RA2/PCs are rarely found in rodent glabrous skin (12), and are located instead in the joints and periosteum (17, 21, 56). Notably, mouse fore and hind paws differ in LTMR innervation density and RA sensitivity (15). Vibrotactile sensitivity is also tuned to fit a species’ particular needs, which can range from tool use to hunting or predator detection (57–59).

Recent work shows that mouse RA2/PCs are sensitive to vibrations into the kHz range and that neurons in primary somatosensory cortex (S1) do not respond with precise spike timing at such high frequencies but, instead, that different frequencies are encoded by the activation of differently tuned neurons (17), like in auditory cortex [(60)); but see ((61))]. Precisely timed spikes have been observed in rat (62) and monkey (42, 54) S1 neurons for stimuli up to 700 or 800 Hz. Cortical neurons spike on intermittent stimulus cycles because of irregular skipping (i.e. they do not entrain 1:1) but remain phase-locked to the stimulus (41). Separate work suggests that rat LTMRs start to desynchronize at frequencies >350 Hz because of reduced entrainment (39), and so encoding kHz vibrations may necessitate a different coding scheme than for the lower vibration frequencies typically studied. Most textures evoke vibrations <400 Hz (63). We did not test >400 Hz for technical reasons, and none of our RAs had remarkably large receptive fields, suggesting we sampled RA1s exclusively. If so, then rodent RA1s can encode frequencies typically ascribed to RA2/PCs in other species (i.e. 100-400 Hz), leaving rodent RA2/PCs to encode even higher frequencies. This implies that rodents encode vibrations up to 400 Hz using heterogeneously tuned RA1s rather than dichotomously tuned RA1s and RA2/PCs. By extension, independent control of S1 spike rate and timing by RA1s and RA2/PCs, respectively, as demonstrated in monkeys (54), would not apply in rodents despite both rate and temporal codes being used in rodent S1 (41, 64, 65).

Consistent with other species, rodent RAs responded transiently to sustained pressure (**Fig. 1A**) but their onset-offset responses were variable. Heterogeneity in RA onset-offset responses was also found in humans and raccoons (48, 66), but not cats (22). These studies agree that RAs favour stimulus onset and, if there is any heterogeneity it typically lies in the offset response (48, 66). Most rodent RAs responded to both stimulus onset and offset (**Fig. 3E**), but based on their differing thresholds, generally preferred either onset or offset (**Fig. 4C**). Recent work in mice has identified two different populations of RAs innervating Meissner corpuscles that respond at stimulus onset and offset (TrkB^+^) or only onset (Ret^+^) (16). Similarly, our models separated into two groups (**Fig. 4C**) but there was no consistent alignment between our ON/OFF groups and the clusters in **Figure 3**. Cluster 1 neurons fire preferentially at onset but some can also fire at offset (albeit with an offset threshold ≥ onset threshold; **Fig. 4C**), which is not consistent with the onset-only spiking exhibited by Ret+ neurons (16), and their slower conduction velocity is actually more consistent with TrkB+ neurons (16). Overall, our results suggest that physiologically defined RA “clusters” do not correspond with molecularly defined Ret^+^ or TrkB^+^ afferents, and that functional heterogeneity is better conceptualized as a continuum.

To explore the basis for RA frequency preferences, we fit GLMs to experimentally measured RA responses (**Fig. 4**). This revealed a continuum of stimulus filter widths (**Fig. 5A**). Narrower filters reflect a higher preferred frequency. A true differentiator has a filter with balanced positive and negative components, whereas an integrator has a broader filter with a positive bias (67). RA stimulus filters were heterogeneous in the balance of their positive and negative components (**Fig. 5C, D**). Stimulus filter shape depends on the specialized end organs with which LTMRs associate (21) as well as the voltage-gated ion channels that control how receptor potentials are converted into spike trains (68); for example, dysfunction of the Kv7.4 channel has been shown to increase RA sensitivity to low-frequency vibration (18). In contrast, the post-spike filter controls firing rate gain as a function of stimulus intensity (69, 70) and limits entrainment as stimulus frequency is increased (**Fig. 7D**). By controlling refractoriness, the post-spike filter affects when a neuron can respond again to a repeating input, whereas the stimulus filter controls the preferred shape (kinetics) of that input. We did not observe any relationship between the stimulus filter and post-spike filter shapes within a given neuron, but our results confirm that rodent RAs exhibit exquisite spike timing precision and can thus support temporal coding.

Broad overlapping tuning curves support combinatorial coding rather than labelled lines (71–74). Like trichromacy, where coactivation of broadly tuned cone photoreceptors supports colour vision, touch appears to rely on the relative activation of broadly tuned LTMRs (which implies that differently tuned LTMRs are co-activated) rather than relying on which particular type of narrowly tuned LTMR is uniquely activated (71–74). Differently tuned LTMRs might fall into distinct groups (as seems to occur in primates) or they might fall along a continuum (as our data suggests for rodents). In either case, sets of heterogeneously tuned RAs convey more information that sets of homogeneously tuned RAs (**Fig. 8**). One might reasonably expect sets of discretely tuned LTMRs based on association with qualitatively different end organs, but heterogeneity in the end organs themselves, in other aspects of the mechanotransduction process (e.g. skin properties), or in LTMR excitability might all contribute to smearing the functional differences between transcriptionally defined afferent types (75). But this does not mean that molecular genetic differences are not important in other ways. For instance, recent work showed that receptive fields of Ret^+^ and TrkB^+^ RAs homotypically tile within subtypes, but heterotypically overlap across subtypes, which is beneficial in terms of spatial acuity (16). Of course, somatosensation extends to thermal, chemical, and noxious mechanical stimuli, which involve other types of afferents not considered in this study.

In summary, we found that rodent RAs respond heterogeneously to vibrotactile stimulation. Further analysis of their tuning curves revealed a continuum of frequency preferences across the RA population, which our modelling suggests is supported by a continuum of stimulus filter widths. Rodent RAs also exhibit heterogeneity in their post-spike filter, which is independent from the heterogeneity in stimulus filter. RAs can modulate their firing rate to support rate coding of stimulus intensity, yet also produce precisely timed (phase-locked) spikes as required for temporal coding of stimulus frequency. In that regard, individual rodent RAs are not so different from their primate counterparts. That said, rodent RAs are arguably more heterogeneous in their tuning, which is not necessarily a good or bad thing, but might have implications for how information is decoded by downstream neurons.

